# Contrasting the form and strength of pre- and postcopulatory sexual selection in a transparent worm with fluorescent sperm

**DOI:** 10.1101/2023.03.22.533851

**Authors:** Lucas Marie-Orleach, Matthew D. Hall, Lukas Schärer

## Abstract

Sexual traits may be selected during multiple consecutive episodes of selection, occurring before, during, or after copulation. The overall strength and shape of selection acting on sexually selected traits may thus be determined by how selection (co-)varies along different episodes. However, it is challenging to measure pre- and postcopulatory phenotypic traits alongside variation in fitness components at each different episode. Here, we used a transgenic line of the transparent flatworm *Macrostomum lignano* expressing green fluorescent protein (GFP) in all cell types, including sperm cells, enabling *in vivo* sperm tracking. We exposed GFP(+) focal worms to three groups in which we assessed their mating success, sperm-transfer efficiency, and sperm fertilising efficiency. Moreover, we measured 13 morphological traits on the focal worms to study the fitness landscape in multivariate trait space. We found linear selection on sperm production rate arising from pre- and postcopulatory components, and on copulatory organ shape arising from sperm fertilising efficiency. We further found nonlinear (mostly concave) selection on combinations of copulatory organ and sperm morphology traits arising mostly from sperm-transfer efficiency and sperm fertilising efficiency. Our study shows that contrasting patterns of phenotypic selection are observed by measuring how sexual selection builds-up over consecutive episodes of selection.

## Introduction

Sexual selection may act before, during, and after copulation, and it represents a compelling framework to explain the evolution of sexual traits involved in mate acquisition, copulation, and fertilisation (Andersson 1994; Birkhead et al. 2009; Prokuda and Roff 2014). Traits under precopulatory sexual selection affect the opportunity to copulate. For instance, in the fungus beetles, *Bolitotherus cornutus*, males with longer horns have better access to females and therefore sire more offspring than their competitors (Conner 1988). Traits under postcopulatory sexual selection affect the number of offspring sired per mating opportunity. For instance, in the cricket *Gryllus bimaculatus*, males producing more and smaller sperm sire more offspring than their competitors (Gage and Morrow 2003). However, pre- and postcopulatory sexual selection may not be independent from one another. For instance, males that have better access to females may consistently sire more (e.g., McDonald et al. 2017; McCullough et al. 2018) or fewer (e.g., De Nardo et al. 2021) offspring per mating opportunity than their competitors. In fact, we know very little about how selection operates across pre- and postcopulatory fitness components, despite precopulatory traits often co-varying with postcopulatory sexual traits, either positively or negatively (reviewed in Mautz et al. 2013; Evans and Garcia-Gonzalez 2016).

Sexual selection studies have traditionally focused on either the interaction between pre- and postcopulatory fitness components (McDonald et al. 2017; McCullough et al. 2018), or on measuring selection for combinations of pre- and postcopulatory traits on total fitness (e.g., Devigili et al. 2015; House et al. 2020). Bridging this knowledge gap instead requires the formal comparison of selection acting on the combinations of phenotypic traits that determine pre- and postcopulatory fitness. If the strength or direction of selection changes across each pre- and postcopulatory fitness components, then the net selection acting on any given trait may be reinforced or negated depending on whether each episode acts synergistically or antagonistically. In considering indirect selection on correlated traits, it is also key to study how trait variation predicts fitness in components where selection is expected occur. For instance, sperm traits may determine male reproductive success, not because they confer higher success in sperm competition, but instead because they are correlated with a trait under precopulatory sexual selection.

Partitioning episodes of sexual selection has long been a feature of sexual selection studies, but it is typically applied to the contrasting effects of male competition and female choice (reviewed in Hunt et al. 2009). Here the relationship between trait variation and relative fitness is used to formally assess the strength of sexual selection, which is typically assessed through the regression coefficient of a linear regression of fitness on one or multiple traits (Lande and Arnold 1983; Arnold and Wade 1984; Jones 2009; Henshaw et al. 2018). This approach has documented strong selection on traits that were assumed to be involved in pre- and postcopulatory processes (reviewed in Simmons and Moore 2009; Prokuda and Roff 2014). Sexual selection may, however, act in more complex and nonlinear ways, including quadratic selection that either favours/opposes intermediate trait values and correlational selection that may act on multiple traits simultaneously (Lande 1979; Lande and Arnold 1983; Henshaw and Zemel 2017). This body of literature has shown that sexual selection often acts on combinations of pre- and/or postcopulatory traits (e.g., Bentsen et al. 2006; Hall et al. 2008; Simmons et al. 2009; Oh and Shaw 2013; Devigili et al. 2015; House et al. 2016, 2020). For instance, in the red flour beetle, *Tribolium castaneum*, male and female genitalia are under concave selection, *i.e.*, intermediate sizes and shapes are favoured by selection (House et al. 2020). Moreover, in the live-bearing fish, *Poecilia reticulata*, selection measured on male reproductive success indicates multivariate fitness landscapes involving both male morphological traits and sperm velocity (Devigili et al. 2015).

To our knowledge, formal comparisons of the strength and form of selection arising from pre- and postcopulatory components of sexual selection have only been obtained once, namely in the broad-horned flour beetle *Gnatocerus cornutus* (House et al. 2016) (albeit in a non-competitive setting). That study found that precopulatory mating success and postcopulatory fertilisation success favoured similar male genital phenotypes, but that concave selection on male body size arose exclusively from precopulatory mating success. More studies along these lines are clearly needed. Arguably, the scarcity of such studies may be due to the (i) high sample sizes required to perform multivariate selection analyses (Green 1991; Simon et al. 2022), and (ii) the challenges in measuring pre- and postcopulatory traits alongside the relevant – and possibly multiple – pre- and postcopulatory fitness components. For instance, although postcopulatory processes are often measured as a single episode of selection, in many species male sperm competitiveness is in fact considered to consist of “(i) the relative number of sperm of different males that enter the fertilising pool; and (ii) the relative fertilisation efficiency of an ejaculate, after controlling for sperm number” (Pizzari and Parker 2009).

Here, taking advantage of powerful features of the free-living flatworm model *Macrostomum lignano* (see below), we studied the strength and form of selection acting on multiple morphological traits with respect to four pre- and postcopulatory fitness components. We measured 13 reproductive morphology traits (including body size, testis size, ovary size, seminal vesicle size, four male copulatory organ traits, and five sperm traits), and we used multivariate selection analyses to statistically test if pre- and postcopulatory sexual selection act synergistically or antagonistically, target similar or dissimilar trait combinations, and differ in the form of selection (directional vs. nonlinear).

Specifically, we sequentially exposed focal worms to three independent mating groups (Figure 1), and assessed, in each group, the individual success of the focal in four fitness components (as defined in Marie-Orleach et al. 2016, 2021). In brief, we decomposed focal male reproductive success into the following four multiplicative fitness components: 1) partner fecundity (*i.e.*, the number of offspring produced by all potential partners through their female sex function), 2) mating success (*i.e.*, the proportion of copulations in which the focal was involved), 3) sperm-transfer efficiency (*i.e.*, the proportion of focal sperm among the sperm received by all potential partners, given the focal mating success) and 4) sperm fertilising efficiency (*i.e.*, the proportion of offspring produced that are sired by the focal, given the proportion of focal sperm received) (see Marie-Orleach et al. 2016, 2021 for more details). We could estimate the last two fitness components because *in vivo* tracking of sperm is feasible in this transparent worm (Janicke et al. 2013; Marie-Orleach et al. 2014; Wudarski et al. 2017). By using transgenic focals that express green fluorescent protein (GFP) in all cell types, including the sperm cells, one can easily distinguish sperm cells donated by GFP(-) wild-type competitors from those that are donated by the GFP(+) focals, directly inside the female reproductive tract of living partners. This provides powerful opportunities to quantify fitness components that are usually difficult to observe, such as the number of sperm cells that a GFP(+) focal individual has successfully transferred to partners, and the resulting number of sired offspring (Janicke et al. 2013; Marie-Orleach et al. 2014, 2016, 2021).

**Figure 1.**
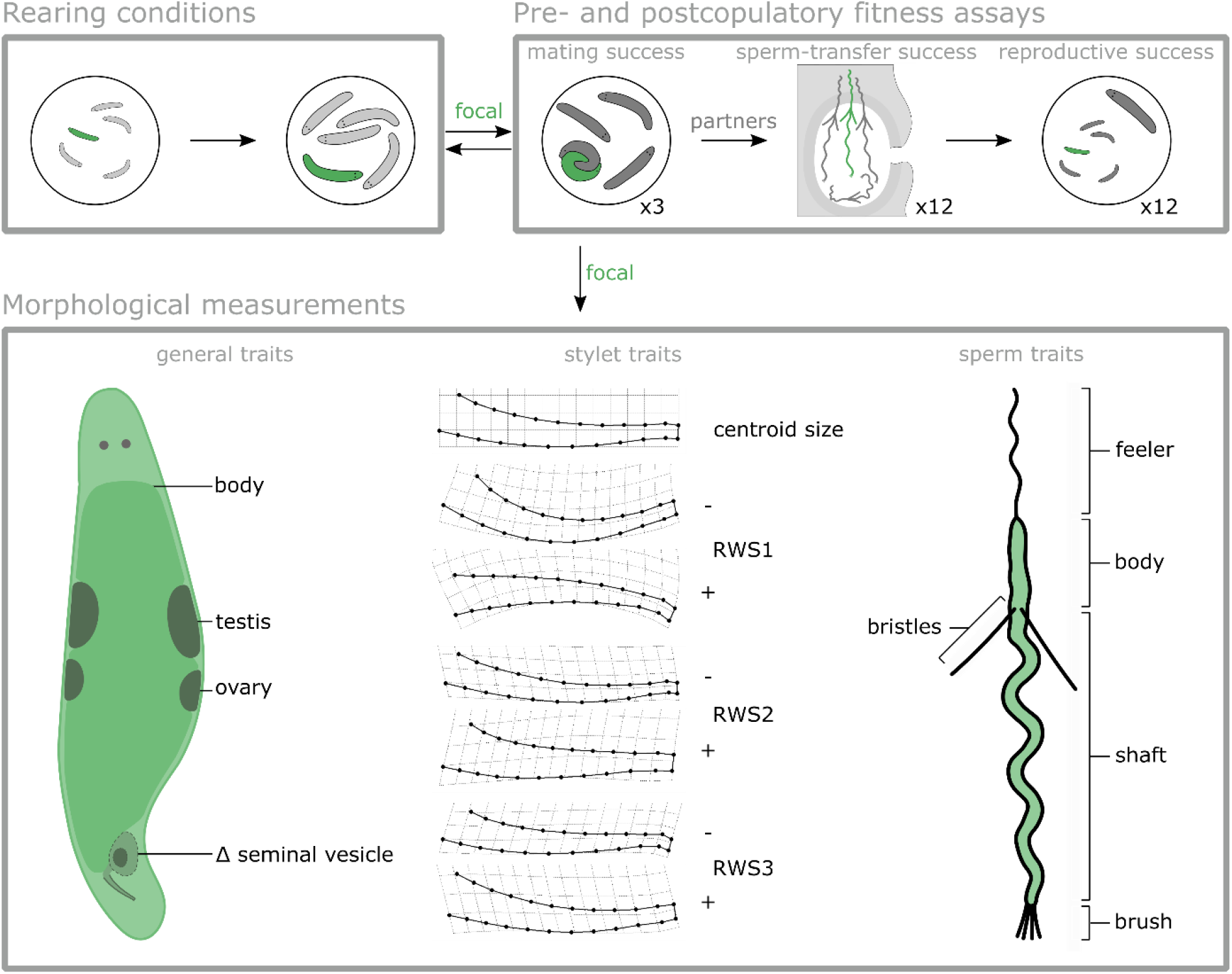
Overview of the experimental design used to measure pre- and postcopulatory morphological traits and fitness components. We allowed GFP(+) focal worms (green) to grow up under ‘Rearing conditions’ with four GFP(-) partners, and then performed ‘Pre- and postcopulatory fitness assays’ by subsequently exposing the focal to three independent groups of four worms to assess mating success, sperm-transfer success and reproductive success, after which we performed ‘Morphological measurements’ on different traits of the focal. We measured 13 traits that were divided into three trait sets. The ‘general traits’ set included the body, testis, ovary, and Δ seminal vesicle sizes (*i.e.*, the change in seminal vesicle size after 2 days of isolation). The ‘stylet traits’ set included the centroid size and the first three relative warp scores of a geometric morphometrics analysis. Diagrams show the configurations of the consensus stylet shape (top), the maximum and minimum values of the first relative warp score (RWS1), which mainly captured the overall stylet curvature, the second relative warp score (RWS2), which mainly captured the width of the stylet, and the third relative warp score (RWS3), which mainly captured the orientation of the tip. RWS1, RWS2 and RWS3 explained 55%, 18% and 12% of the variation in stylet shape, respectively. The ‘sperm traits’ set included the feeler, body, shaft, brush and bristles sizes. See Methods for details.

Furthermore, we collected data to characterise selection acting on trait combinations that are potentially important during pre- and postcopulatory selection (Figure 1). These traits include testis size, which has previously been found to predict mating success (Janicke and Schärer 2009a,b), sperm production rate (Schärer and Vizoso 2007), sperm-transfer success (Janicke and Schärer 2009a; Marie-Orleach et al. 2016), and male reproductive success in this worm (Marie-Orleach et al. 2016; Vellnow et al. 2018). We also measured body size, ovary size, Δ seminal vesicle size (*i.e.*, the increase in seminal vesicle size of focals after two days in isolation, a measure of the sperm production rate), the size and shape of the male copulatory organ (the so-called stylet) using geometric morphometrics, and five different sperm morphology traits to test how these predict male fitness (Janicke and Schärer 2009a, 2010; Marie-Orleach et al. 2016).

Intense postcopulatory sexual selection is expected in simultaneous hermaphrodites (Charnov 1979; Schärer and Pen 2013; Schärer et al. 2014; Marie-Orleach et al. 2021), and we expected testis size and Δ seminal vesicle size to be under directional positive selection, since both may positively contribute to higher success in sperm-transfer efficiency. In contrast, a trade-off between the male and female sex functions may lead to negative correlations between male reproductive success and traits involved in the female sex function, such as ovary size (Anthes et al. 2010). Several different hypotheses have been proposed to explain the evolution of genitalia. Because interspecific studies usually find a lower allometric relationship for animal genitalia size than for other organs (*i.e.*, the one-size-fits-all hypothesis), animal genitalia are often thought to be under concave selection (Eberhard et al. 1998). Arnqvist (1997) further predicts that the pleiotropism, the lock-and-key, and the sexual selection hypotheses should, respectively, lead to no selection, concave selection, and directional selection within species. Regardless of the type of selection, we expected selection on stylet traits to arise mostly from sperm-transfer efficiency. And finally, we expected directional selection on sperm traits arising from the sperm fertilising efficiency fitness component. In particular, *M. lignano* displays a complex sperm morphology (Figure 1), including lateral stiff bristles that may be important for the outcome of sperm competition (Vizoso et al. 2010; Schärer et al. 2011).

## Material and Methods

The data of the present study are associated with an experiment that is described in considerable detail elsewhere (Marie-Orleach et al. 2021). In the previous paper we reported analyses using a variance-based approach that did not consider the morphology of the focal worms (Marie-Orleach et al. 2021), and here we report the results of a trait-based approach to studying sexual selection that includes morphological data on the focal individuals. In the following, we briefly describe the parts of the material and methods shared with Marie-Orleach et al. (2021), and we also more fully explain the parts that are specific to the present study.

### Model organism

*Macrostomum lignano* (Macrostomorpha, Platyhelminthes) is a free-living flatworm found in the upper intertidal zone of the Mediterranean Sea (Ladurner et al. 2005; Schärer et al. 2020), and it can be easily cultured in the laboratory (Wudarski et al. 2020). The worms are simultaneous hermaphrodites, obligately outcrossing (Schärer and Ladurner 2003), and highly promiscuous (Schärer et al. 2004; Janicke and Schärer 2009a). The copulation is reciprocal, and consists of the intromission of the male copulatory organ (called stylet) into the female sperm-receiving and sperm-storage organ (called female antrum) of the partner (Schärer et al. 2004; Vizoso et al. 2010). Because the worms are highly transparent, it is possible to observe and measure internal structures *in vivo*, such as testis and ovary size (Schärer and Ladurner 2003; Janicke et al. 2013), stylet morphology (Janicke and Schärer 2009a; Marie-Orleach et al. 2016), and the number of sperm cells stored inside the female antrum (Janicke et al. 2011; Marie-Orleach et al. 2016, 2021). Moreover, sperm traits can be measured after amputation of a worm’s tail plate (Janicke and Schärer 2010).

*In vivo* sperm tracking is possible in *M. lignano* thanks to transgenic lines expressing GFP (Janicke et al. 2013; Marie-Orleach et al. 2014, 2021; Wudarski et al. 2017). Moreover, one can use the GFP marker to assign parentage in offspring (Marie-Orleach et al. 2014, 2021). Importantly, sexual behaviour and reproductive performances of GFP(+) individuals are not different from those of GFP(-) individuals (Marie-Orleach et al. 2014). In this study, we used two outbred cultures, the GFP(+) BAS1 (Marie-Orleach et al. 2016; Vellnow et al. 2018) and the GFP(-) LS1 (Marie-Orleach et al. 2013). These cultures are expected to be genetically similar because BAS1 was established by introgression of the GFP marker into the LS1 culture (Marie-Orleach et al. 2016; Vellnow et al. 2018).

### Experimental set-up

Our experimental set-up allowed us to measure the mating success, sperm-transfer success and reproductive success of focal worms, for which we also measured 13 morphological traits (see below and Figure 1). For logistic reasons, the biological replicates used in the experiment were split into eight batches, treated sequentially (three to six days apart), but in the following reporting we set day 1 as the first day for each batch.

#### Rearing conditions

To obtain same-aged individuals we, on day 1, allowed adult worms of the GFP(+) and GFP(-) cultures to lay eggs in Petri dishes for 24h and, on day 6, we placed the resulting offspring in 24-well tissue culture plates to create 20 biological replicates per batch. Each biological replicate included (i) one group made up of one GFP(+) focal and four GFP(-) partners (called the A groups), and (ii) three groups made up of four GFP(-) individuals (called the B, C, and D groups) that we used as partners of the focal (Figure 1). We then transferred all groups to new wells with fresh algae every 6 to 10 days. Importantly, in order to measure selection in individuals that had reached a steady state of sperm production, mating activity, sperm receipt, and egg production, we kept the worms in their A groups for several days after they had reached sexual maturity.

#### Mating success

We then estimated the focal’s mating success on day 25 to 30 as explained in Marie-Orleach et al. (2021). In brief, we placed each focal in a mating chamber together with the four worms of its B group. To visually distinguish the focal from the partners, we placed all members of the A group into a well containing a blue vital dye for the 24h before the mating trial (which does not affect sexual performance; Marie-Orleach et al. 2013). We then placed the five worms in an 8 µL drop of artificial sea water between two microscope slides and video recorded all their interactions for 3 h. In total, we gathered 1,350h of copulation interactions (analysed blindly with regards to replicate identity), which contained 26,203 copulations. For each mating group we counted the total number of copulations (total matings) and the number of copulations in which the focal was involved (focal matings). After the mating trials, we transferred the focal back into its A group, and isolated all four members of the B group to assess the sperm-transfer success of the focal during the mating trial.

#### Sperm-transfer success

We estimated the proportion of sperm cells received from the focal by its four partners following the protocol explained in Marie-Orleach et al. (2021). In brief, we recorded movies of the female antrum of each partner, first under bright-field illumination and then under epifluorescence illumination to, respectively, assess (i) the total number of sperm cells in all four potential partners (total sperm) and (ii) the number of these sperm that were GFP(+) (focal sperm). The 3,600 resulting antrum movies were analysed blindly with regards to replicate identity. The procedure used provides highly repeatable sperm counts (Marie-Orleach et al. 2014). Note that, when we could not assess total sperm in worms because they had eggs in their female antrum (731/1800), we used the average total sperm counts computed from worms in which this could be estimated (*i.e.*, 21 sperm cells).

#### Reproductive success

We estimated the reproductive success of the focal by letting the partners lay eggs in isolation for 12 days, as explained in Marie-Orleach et al. (2021) – yielding 10,452 offspring in total. We counted the total number of offspring produced across all four potential partners (total offspring) and determined the number of these that were GFP(+) (focal offspring). We computed the proportion of offspring sired by the focal by dividing the sum of focal offspring by the sum of number of total offspring.

On the two subsequent days, we repeated all of the above-mentioned steps (*mating success*, *sperm-transfer success*, and *reproductive success*) by placing each focal together with the four individuals of their C and D groups, respectively. We thus measured the same pre- and postcopulatory components of male reproductive success for each focal in three independent mating groups of four worms, sampled from the same pool of same-aged outbred worms.

#### Morphological measurements

On the next day, we assessed a number of morphological traits of the focals by following protocols described elsewhere (Schärer and Ladurner 2003; Janicke and Schärer 2009a). Briefly, these protocols consist of anaesthetising and squeezing the worm in between a microscope slide and a haemocytometer cover slip in a standardized way and taking digital pictures of the entire body, the two testes, the two ovaries, the seminal vesicle, and the stylet. For this, we used a Leica DM2500 microscope, an Imaging Source DFK 41AF02 camera, and BTV Pro 6.0b7. Focals were then isolated for 2 days, after which we measured them again to assess the morphological traits a second time, in order to i) reduce measurement error and ii) estimate the increase in seminal vesicle size over 2 days (i.e., Δ seminal vesicle size), which can be used as a reliable proxy for the sperm production rate (Schärer and Vizoso 2007). We analysed the pictures blindly with regards to replicate identity, using ImageJ, and we averaged the two measurements to assess the body size, testis size and ovary size. Moreover, we assessed stylet morphology by using a geometric morphometric approach as in Janicke and Schärer (2009a), where we first analysed all 320 stylet pictures in a single analysis, and then, for each focal, averaged the two measurements of stylet centroid size (CS) and of the first three relative warp scores (RWS). These RWS values together explained 85% of the total variance observed in stylet shape, and mainly captured the overall curvature of the stylet (RWS 1), the width of the stylet (RWS2), and the orientation of the tip of the stylet (RWS 3). See Figure 1 for visualisations.

Immediately after the second measurement, we assessed five sperm traits following an established protocol (Janicke and Schärer 2010), by amputating, squeezing, and thus rupturing the tail plate of the worm in a tiny drop, so that the sperm cells are released into the medium and become accessible for imaging. The pictures were then analysed using ImageJ, blindly with regards to replicate identity, to assess the length of the feeler, body, shaft, bristles and brush (Janicke and Schärer 2010). Note that we measured both bristles and used the averaged length. We assessed on average 7.7 sperm cells per individual (range from 1 to 11), and used the averaged values for the statistical analysis.

#### Penetrance of the GFP marker

Finally, as explained in Marie-Orleach et al. (2021), we estimated the penetrance of the GFP marker for each focal worm by pairing them with a virgin GFP(-) individual. We then assessed the GFP status of the resulting offspring (47.7 offspring screened per focal on average), which led to the exclusion of two focals. One produced 47% GFP(+) offspring, and the other one produced no offspring with the virgin GFP(-) individual. The other focals produced either 100% (n=135) or between 90% and 100% GFP(+) offspring (n=23), which will have little influence on our estimates of male reproductive success.

### Data analysis

To compare the strength and form of selection acting on the measured morphological traits at pre- and postcopulatory fitness components, we measured (1) linear and (2) nonlinear selection on the original morphological traits, and (3) the nonlinear selection on composite traits that are more suited to test for nonlinear selection. Before the analysis, all fitness data were relativised to a mean of 1 (Jones 2009). Also, we transformed testis size (square root) and ovary size (log_10_) to account for data skewness, and then we standardised all morphological traits to a mean of 0 and a standard deviation of 1 (Jones 2009). Importantly, we split our 13 morphological traits into three sets of traits. Such a procedure was necessary to avoid overfitting issues, which, in our case, would lead to poor power to detect nonlinear selection on the original morphological traits, and spuriously significant nonlinear effects on the composite traits. Given our sample size, we restricted the number of traits to a maximum of five (which leads to 20 total predictors, including the quadratic and cross-product two-way interactions) (Green 1991). The traits were split as follows: the *general traits set* (body size, testis size, ovary size, and Δ seminal vesicle size), the *stylet traits set* (CS, RWS1, RWS2 and RWS3), and the *sperm traits set* (feeler size, body size, shaft size, bristle size and brush size). We split the 13 traits in this way due to *a priori* expectations. Namely, we expected traits of the *general traits set* to be involved mainly in *mating success* and *sperm-transfer efficiency*, those of the *stylet trait set* in *sperm-transfer efficiency*, and those of the *sperm traits set* in *sperm fertilising efficiency*.

Our initial sample size was 160 replicates, but because we lost replicates due to developmental errors (n=8), the penetrance of the GFP marker (n=2), and missing measurements of one or more traits (n=11), our final sample size was reduced to 139 replicates.

#### Selection on the original trait sets

We first studied whether there was linear and nonlinear selection acting over all traits contained in a given trait set. For this, we used a sequential model building approach that statistically compares the fits of models with and without the terms of interest (see appendix A of Chenoweth and Blows 2005). Here, we specifically tested if the fits of linear models predicting fitness, and including only the intercept, are improved by adding, first, the linear terms of all traits contained in a trait set (*i.e.*, testing for linear selection), and then, all quadratic and cross-product terms (*i.e.*, testing for nonlinear selection). We did this analysis separately for all three trait sets (*i.e.*, general, stylet, and sperm traits sets), and on relative male reproductive success and the four fitness components (*i.e.*, *mRS*\*, *F*\*, *MS*\*, *STE*\*, *SFE*\*).

Second, to test if linear and nonlinear selection measured in all three trait sets were consistent across the four fitness components, we used again the sequential model building approach (Chenoweth and Blows 2005). We built linear models predicting fitness in all four fitness components together, and including the intercept and the linear terms, and tested if their fits were statistically improved by the addition of the interaction term *linear term × fitness component* (Chenoweth and Blows 2005). Significant interaction terms indicate that linear selection is different across fitness components (Chenoweth and Blows 2005; House et al. 2016). We further tested if patterns of nonlinear selection were consistent over the fitness components by testing if the addition of the interaction terms *nonlinear terms × fitness component* statistically improved the fit of the models including the nonlinear terms (Chenoweth and Blows 2005). When selection was inconsistent across fitness components, we tested if selection was different between each pair of fitness component<u>s</u>. Note that because these analyses required using each replicate several times in a single analysis (i.e., once for each fitness component), we performed additional analyses including replicate IDs as a random effect, which did not qualitatively change the outcomes (data not shown).

We then estimated the linear selection gradients (*β*) for each trait, computed through the partial regression coefficients of a multiple linear regression of *mRS*\*, *F*\*, *MS*\*, *STE*\*, and *SFE*\* separately on all morphological traits (Lande and Arnold 1983), which indicated which specific morphological traits were experiencing linear selection. We also computed the matrix containing the quadratic and cross-product selection gradients, called the γ matrix, which we estimated through a multiple linear regression including all quadratic (γii), and cross-product terms (γ_ij_). We doubled the quadratic regression coefficients so that the estimates for concave and convex forms of selection correspond to the Lande and Arnold (1983) formulation (Stinchcombe et al. 2008). The γ matrix allowed us to determine which specific interaction terms were responsible for the nonlinear effect.

#### Nonlinear selection on composite traits

We studied nonlinear selection by using composite traits describing morphological trait variation, which were generated by canonical rotations of the γ matrix (Phillips and Arnold 1989; Blows and Brooks 2003; Blows 2007). The canonical rotation of the γ matrix generates the so-called **M** matrix, which contains (i) the eigenvectors (*m*_i_) describing the major axes of nonlinear selection (i.e., the loading of each original morphological trait), and (ii) their eigenvalues (λ_i_) describing the strength and form of nonlinear selection. A negative eigenvalue indicates a concave relationship between the fitness and the eigenvector (i.e., a “peak” in which high fitness is reached for intermediate values along the eigenvector) whereas positive eigenvalues indicate convex relationship (i.e., a “bowl” in which high fitness is reached for low and high values along the eigenvector). An advantage of the canonical analysis is that the composite traits of variation are orthogonal to one another, which makes the cross-product terms (γ_ij_) null by definition. All nonlinear selection is thus captured by the quadratic terms (γ_ii_), which provides better power to detect nonlinear selection (Blows and Brooks 2003).

We then tested if there was nonlinear selection in each trait set, and if it was consistent over the four fitness components, this time using the composite traits in a sequential model building approach (Chenoweth and Blows 2005; Garant et al. 2007; Parker et al. 2011). As above, we first tested if the additions of the quadratic terms significantly improved the fits of models predicting fitness and including only the linear terms. We tested if nonlinear selection was consistent across the four fitness components by testing if the addition of the interaction terms *nonlinear terms × fitness component* significantly improved the fit of models predicting fitness on all fitness components (Garant et al. 2007; Parker et al. 2011). As above, we ran additional analyses including replicate ID as a random effect, which again did not qualitatively change the outcomes (data not shown).

We then explored the eigenvalues and eigenvectors of the **M** matrix to determine whether the nonlinear selection was concave or convex, and which traits generated nonlinear selection. Statistical significance of the eigenvalues was tested by permutation of the fitness values (10,000 iterations) (Keagy et al. 2016).

## Results

### General traits set

We found significant linear selection in the *general traits set* on *mRS\** (Table 1A), which was due to a significant positive selection gradient of Δ seminal vesicle size on *mRS*\* (Table 2A; Figure 2A). Focals that replenished their seminal vesicle more quickly sired more offspring. Linear selection was not significantly different across the four fitness components (Table S1A), which suggest that the effect of Δ seminal vesicle size on *mRS*\* arises from weak positive selection on multiple fitness components (*MS*\*, *STE*\*, *SFE*\*) (although significantly so only in *MS*\*; Table 2).

**Figure 2.**
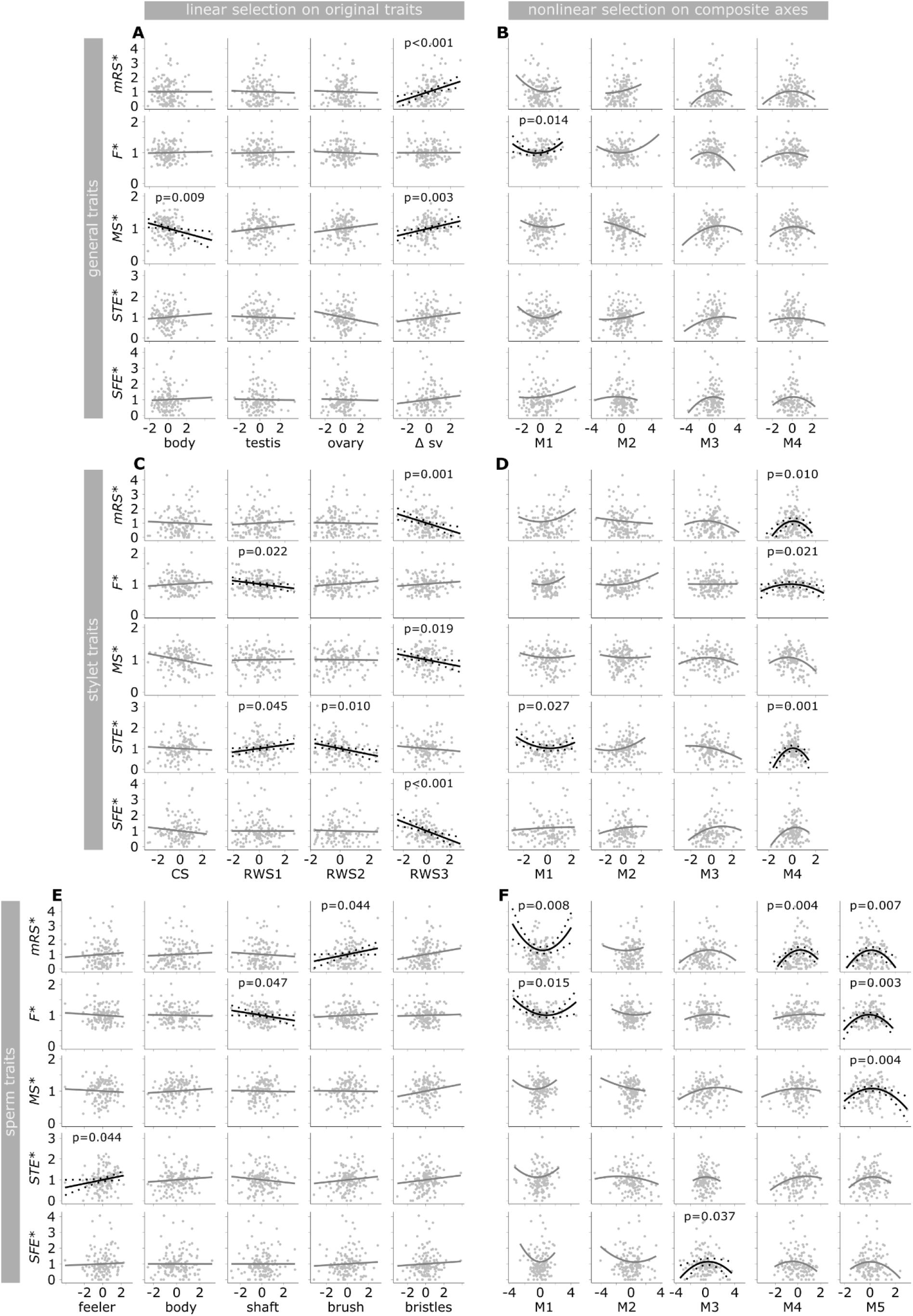
Linear selection (panels A, C and E) and multivariate selection (panels B, D and F) on the 13 morphological traits and the derived composite traits on male reproductive success, and the four fitness components. Grey points represent observed values. Solid lines represent predicted marginal effects of the x-variable from models including either the linear effects of all traits of a trait set (morphological traits, see Table 2A for statistics), or the linear and quadratic effects of all composite traits of a trait set (composite traits, see Table 2B for statistics). Dotted lines represent the 95% confidence intervals. Fit lines are drawn in black when we found *P* values below 0.05, and in grey otherwise. Morphological and composite traits are standardised, and male relative success is relative (see Methods).

**Table 1.**
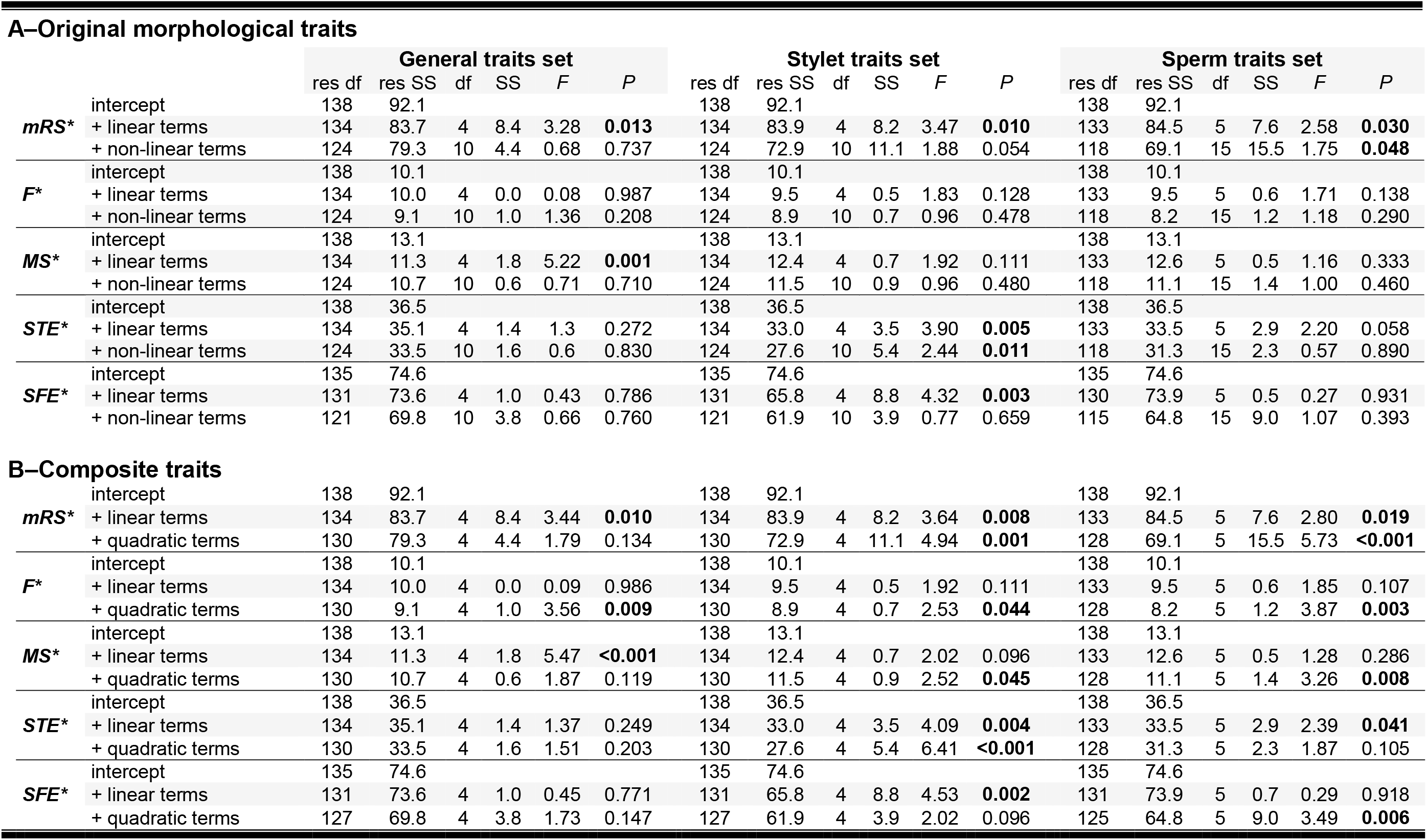
ANOVAs for the full second-order polynomial regressions of male reproductive success (mRS*) and four male fitness components, partner fecundity (F*), mating success (MS*), sperm transfer efficiency (SFE*), and sperm fertilising efficiency (SFE*), on the original morphological traits (A) and composite traits (B) of the general traits, stylet traits, and sperm traits. In the original morphological traits, non-linear terms includes both the quadratic and cross-product terms.

A–Original morphological traits
|  |  | General traits set |  |  |  |  |  | Stylet traits set |  |  |  |  |  | Sperm traits set |  |  |  |  |  |
| --- | --- | --- | --- | --- | --- | --- | --- | --- | --- | --- | --- | --- | --- | --- | --- | --- | --- | --- | --- |
|  |  | res df | res SS | df | SS | F | P | res df | res SS | df | SS | F | P | res df | res SS | df | SS | F | P |
| $mRS^*$ | intercept | 138 | 92.1 | | | | | 138 | 92.1 | | | | | 138 | 92.1 | | | | |
|  | + linear terms | 134 | 83.7 | 4 | 8.4 | 3.28 | **0.013** | 134 | 83.9 | 4 | 8.2 | 3.47 | **0.010** | 133 | 84.5 | 5 | 7.6 | 2.58 | **0.030** |
|  | + non-linear terms | 124 | 79.3 | 10 | 4.4 | 0.68 | 0.737 | 124 | 72.9 | 10 | 11.1 | 1.88 | 0.054 | 118 | 69.1 | 15 | 15.5 | 1.75 | **0.048** |
| $F^*$ | intercept | 138 | 10.1 | | | | | 138 | 10.1 | | | | | 138 | 10.1 | | | | |
|  | + linear terms | 134 | 10.0 | 4 | 0.0 | 0.08 | 0.987 | 134 | 9.5 | 4 | 0.5 | 1.83 | 0.128 | 133 | 9.5 | 5 | 0.6 | 1.71 | 0.138 |
|  | + non-linear terms | 124 | 9.1 | 10 | 1.0 | 1.36 | 0.208 | 124 | 8.9 | 10 | 0.7 | 0.96 | 0.478 | 118 | 8.2 | 15 | 1.2 | 1.18 | 0.290 |
| $MS^*$ | intercept | 138 | 13.1 | | | | | 138 | 13.1 | | | | | 138 | 13.1 | | | | |
|  | + linear terms | 134 | 11.3 | 4 | 1.8 | 5.22 | **0.001** | 134 | 12.4 | 4 | 0.7 | 1.92 | 0.111 | 133 | 12.6 | 5 | 0.5 | 1.16 | 0.333 |
|  | + non-linear terms | 124 | 10.7 | 10 | 0.6 | 0.71 | 0.710 | 124 | 11.5 | 10 | 0.9 | 0.96 | 0.480 | 118 | 11.1 | 15 | 1.4 | 1.00 | 0.460 |
| $STE^*$ | intercept | 138 | 36.5 | | | | | 138 | 36.5 | | | | | 138 | 36.5 | | | | |
|  | + linear terms | 134 | 35.1 | 4 | 1.4 | 1.3 | 0.272 | 134 | 33.0 | 4 | 3.5 | 3.90 | **0.005** | 133 | 33.5 | 5 | 2.9 | 2.20 | 0.058 |
|  | + non-linear terms | 124 | 33.5 | 10 | 1.6 | 0.6 | 0.830 | 124 | 27.6 | 10 | 5.4 | 2.44 | **0.011** | 118 | 31.3 | 15 | 2.3 | 0.57 | 0.890 |
| $SFE^*$ | intercept | 135 | 74.6 | | | | | 135 | 74.6 | | | | | 135 | 74.6 | | | | |
|  | + linear terms | 131 | 73.6 | 4 | 1.0 | 0.43 | 0.786 | 131 | 65.8 | 4 | 8.8 | 4.32 | **0.003** | 130 | 73.9 | 5 | 0.5 | 0.27 | 0.931 |
|  | + non-linear terms | 121 | 69.8 | 10 | 3.8 | 0.66 | 0.760 | 121 | 61.9 | 10 | 3.9 | 0.77 | 0.659 | 115 | 64.8 | 15 | 9.0 | 1.07 | 0.393 |
| B–Composite traits |  |  |  |  |  |  |  |  |  |  |  |  |  |  |  |  |  |  |  |
| $mRS^*$ | intercept | 138 | 92.1 | | | | | 138 | 92.1 | | | | | 138 | 92.1 | | | | |
|  | + linear terms | 134 | 83.7 | 4 | 8.4 | 3.44 | **0.010** | 134 | 83.9 | 4 | 8.2 | 3.64 | **0.008** | 133 | 84.5 | 5 | 7.6 | 2.80 | **0.019** |
|  | + quadratic terms | 130 | 79.3 | 4 | 4.4 | 1.79 | 0.134 | 130 | 72.9 | 4 | 11.1 | 4.94 | **0.001** | 128 | 69.1 | 5 | 15.5 | 5.73 | **<0.001** |
| $F^*$ | intercept | 138 | 10.1 | | | | | 138 | 10.1 | | | | | 138 | 10.1 | | | | |
|  | + linear terms | 134 | 10.0 | 4 | 0.0 | 0.09 | 0.986 | 134 | 9.5 | 4 | 0.5 | 1.92 | 0.111 | 133 | 9.5 | 5 | 0.6 | 1.85 | 0.107 |
|  | + quadratic terms | 130 | 9.1 | 4 | 1.0 | 3.56 | **0.009** | 130 | 8.9 | 4 | 0.7 | 2.53 | **0.044** | 128 | 8.2 | 5 | 1.2 | 3.87 | **0.003** |
| $MS^*$ | intercept | 138 | 13.1 | | | | | 138 | 13.1 | | | | | 138 | 13.1 | | | | |
|  | + linear terms | 134 | 11.3 | 4 | 1.8 | 5.47 | **<0.001** | 134 | 12.4 | 4 | 0.7 | 2.02 | 0.096 | 133 | 12.6 | 5 | 0.5 | 1.28 | 0.286 |
|  | + quadratic terms | 130 | 10.7 | 4 | 0.6 | 1.87 | 0.119 | 130 | 11.5 | 4 | 0.9 | 2.52 | **0.045** | 128 | 11.1 | 5 | 1.4 | 3.26 | **0.008** |
| $STE^*$ | intercept | 138 | 36.5 | | | | | 138 | 36.5 | | | | | 138 | 36.5 | | | | |
|  | + linear terms | 134 | 35.1 | 4 | 1.4 | 1.37 | 0.249 | 134 | 33.0 | 4 | 3.5 | 4.09 | **0.004** | 133 | 33.5 | 5 | 2.9 | 2.39 | **0.041** |
|  | + quadratic terms | 130 | 33.5 | 4 | 1.6 | 1.51 | 0.203 | 130 | 27.6 | 4 | 5.4 | 6.41 | **<0.001** | 128 | 31.3 | 5 | 2.3 | 1.87 | 0.105 |
| $SFE^*$ | intercept | 135 | 74.6 | | | | | 135 | 74.6 | | | | | 135 | 74.6 | | | | |
|  | + linear terms | 131 | 73.6 | 4 | 1.0 | 0.45 | 0.771 | 131 | 65.8 | 4 | 8.8 | 4.53 | **0.002** | 131 | 73.9 | 5 | 0.7 | 0.29 | 0.918 |
|  | + quadratic terms | 127 | 69.8 | 4 | 3.8 | 1.73 | 0.147 | 127 | 61.9 | 4 | 3.9 | 2.02 | 0.096 | 125 | 64.8 | 5 | 9.0 | 3.49 | **0.006** |

**Table 2.**
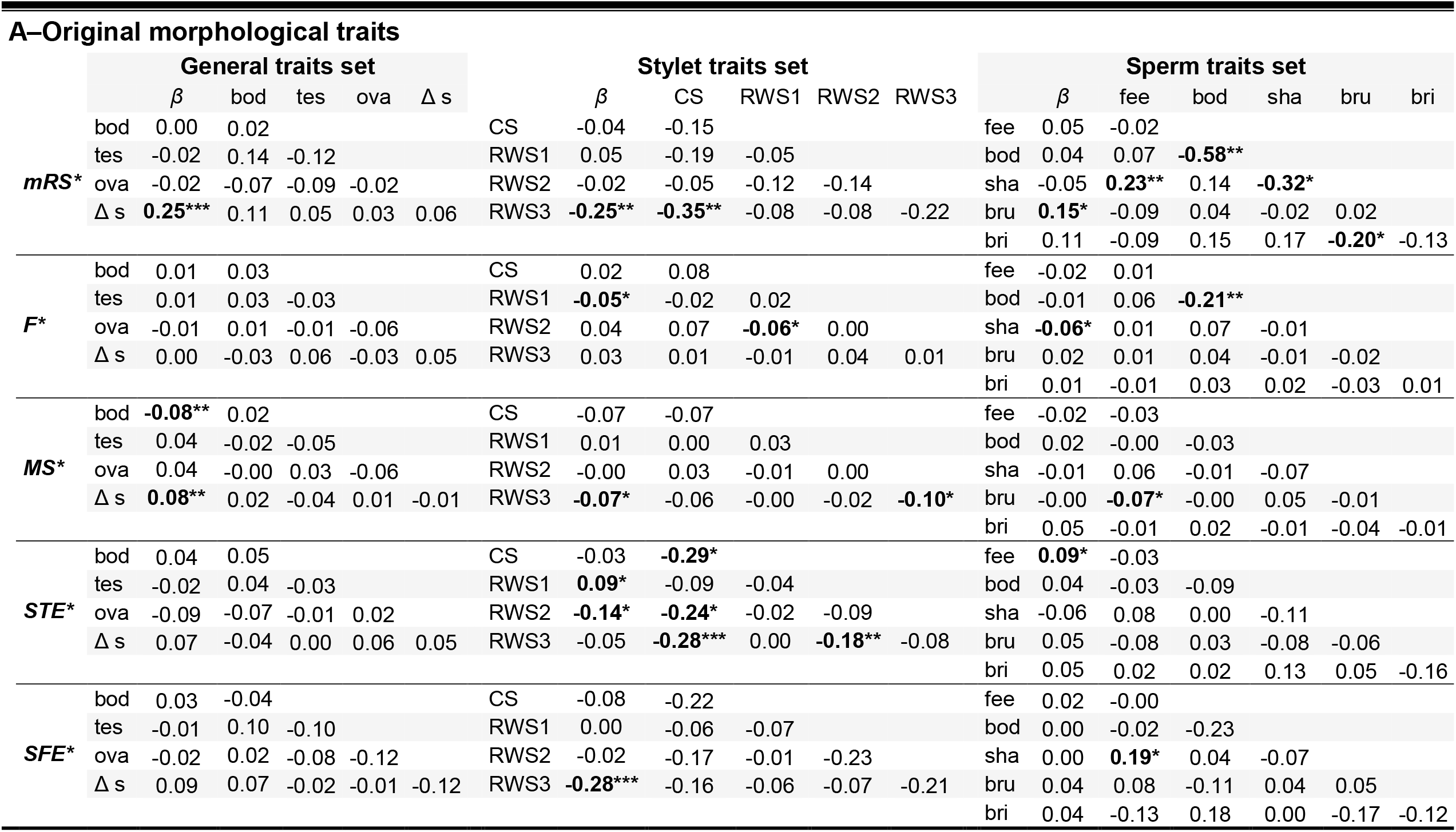
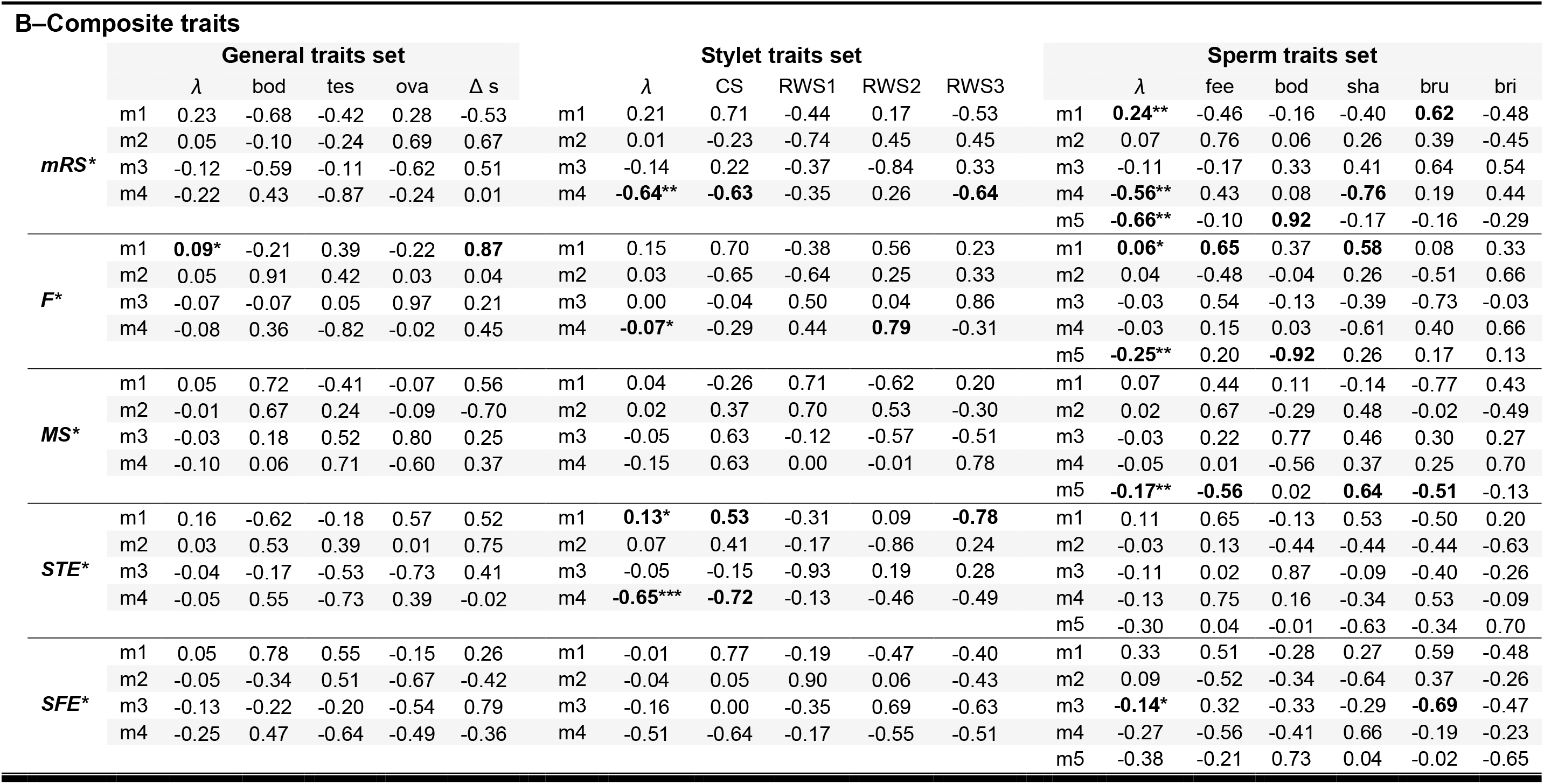
Linear and non-linear selection on each trait set, measured on male reproductive success (mRS*) and on the four male fitness components, partner fecundity (F*), mating success (MS*), sperm transfer efficiency (SFE*), and sperm fertilising efficiency (SFE*). (A) shows the standardised linear selection gradients (β), and the γ matrices showing the standardised quadratic and correlational selection gradients. (B) shows the eigenvalues (λ), and the M matrices of the eigenvectors from the canonical rotation of the γ matrices. Traits are body size (bod), testis size (tes), ovary size (ova), Δ s (Δ seminal vesicle size); stylet’s centroide size (CS), 1^st^ 2^nd^ and 3^rd^ relative warp scores (RWS1, RWS2, RWS3); sperm feeler size (fee), body size (bod), shaft size (sha), brush size (bru) and bristle size (bri). Bold font stands for significant gradients and eigenvalues, and for loadings higher than |0.50| on canonical axes with significant eigenvalues. \**P*<0.05; \*\**P*<0.01; \*\*\**P*<0.001.

We found no evidence of nonlinear selection in the general trait set on *mRS*\* using either the original morphological traits (Table 1A) or the composite traits (Table 1B). Moreover, nonlinear selection did not seem to be different across fitness components (Table S1). These results suggest that there is no nonlinear selection acting on body size, testis size, ovary size, and Δ seminal vesicle size, nor are there any interactions between these traits on *mRS*\*.

### Stylet traits set

We found linear selection in the *stylet traits set* on *mRS*\* (Table 1A), which arose due to a negative effect of stylet RWS3 on *mRS*\* (Table 2A; Figure 2C). This suggested that worms with more bent stylet tips sired more offspring. Patterns of linear selection were different across fitness components (Table S1A). Specifically, we found different patterns of linear selection on *STE*\* and *SFE*\*, and no significant linear selection on *F\** and *MS\** (Table 1A). Interestingly, *STE*\* induced positive linear selection on RWS1 and negative linear selection on RWS2, whereas *SFE*\* induced negative linear selection on RWS3 (Table 2A; Figure 2C). These results suggest that different aspects of stylet morphology are selected on the two postcopulatory fitness components, *STE*\* and *SFE*\*. Specifically, worms with stylets curved away from the seminal vesicle (i.e., high RWS1 values) and narrow stylets (*i.e.*, low RWS2 values) had a selective advantage with respect to *STE*\*, whereas worms with stylets with more bent tips (i.e., low RWS3 values) had a selective advantage with respect to *SFE*\*.

We found evidence for nonlinear selection acting on *mRS*\*, which was almost significant using the original morphological traits (Table 1A), and highly significant using the composite traits (Table 1B). This effect was due to an interaction between centroid size and RWS3 (Table 2A), both of which had negative weights on m4, a composite trait capturing concave selection in *mRS*\* (Table 2B). High *mRS*\* is thus achieved with intermediate m4 values, which may result from either intermediate centroid size and RWS3 values, or from centroid size and RWS3 values that counteract each other, such as stylets that are small and have a straight tip, or that are large and have a more bent tip.

This nonlinear selection was not consistent across fitness components (Table S1B), which was due to differences in *F*\*, *MS*\*, and *STE*\* (Table S3B). We detected nonlinear selection in all these three fitness components (Table 1B), though nonlinear effects were larger in *STE*\* compared to *F*\* and *MS*\* (Table 2B; Figure 2D). Concave and convex nonlinear selection was found in *STE*\* (Table 2B; Figure 3). The composite trait with the largest |eigenvalue| was m4. It displayed concave selection and was mostly affected by centroid size (and RWS2 and RWS3 to a lesser extent), all with negative loadings (orange labels), and thus probably underlay the nonlinear effect we detected on *mRS*\*. The second significant composite trait was m1, which displayed convex selection and was influenced by centroid size and RWS3 but, importantly, with a positive (blue labels) and a negative (orange labels) loading, respectively (Figure 3). This result means that worms with stylets that are small and have straight tips, or are large and have highly bent tips, had higher *STE*\* values. Although this may appear counterintuitive at first sight, the convex effect found on *STE*\* may concord with the concave effect found on *mRS*\*. This is because opposite values in centroid size and RWS3 (i.e., high values in centroid size and low value in RWS, or the other way around) lead to extreme values on m1 on *STE*\*, and to average values on m4 on *mRS*\*, which are both associated with high fitness values.

**Figure 3.**
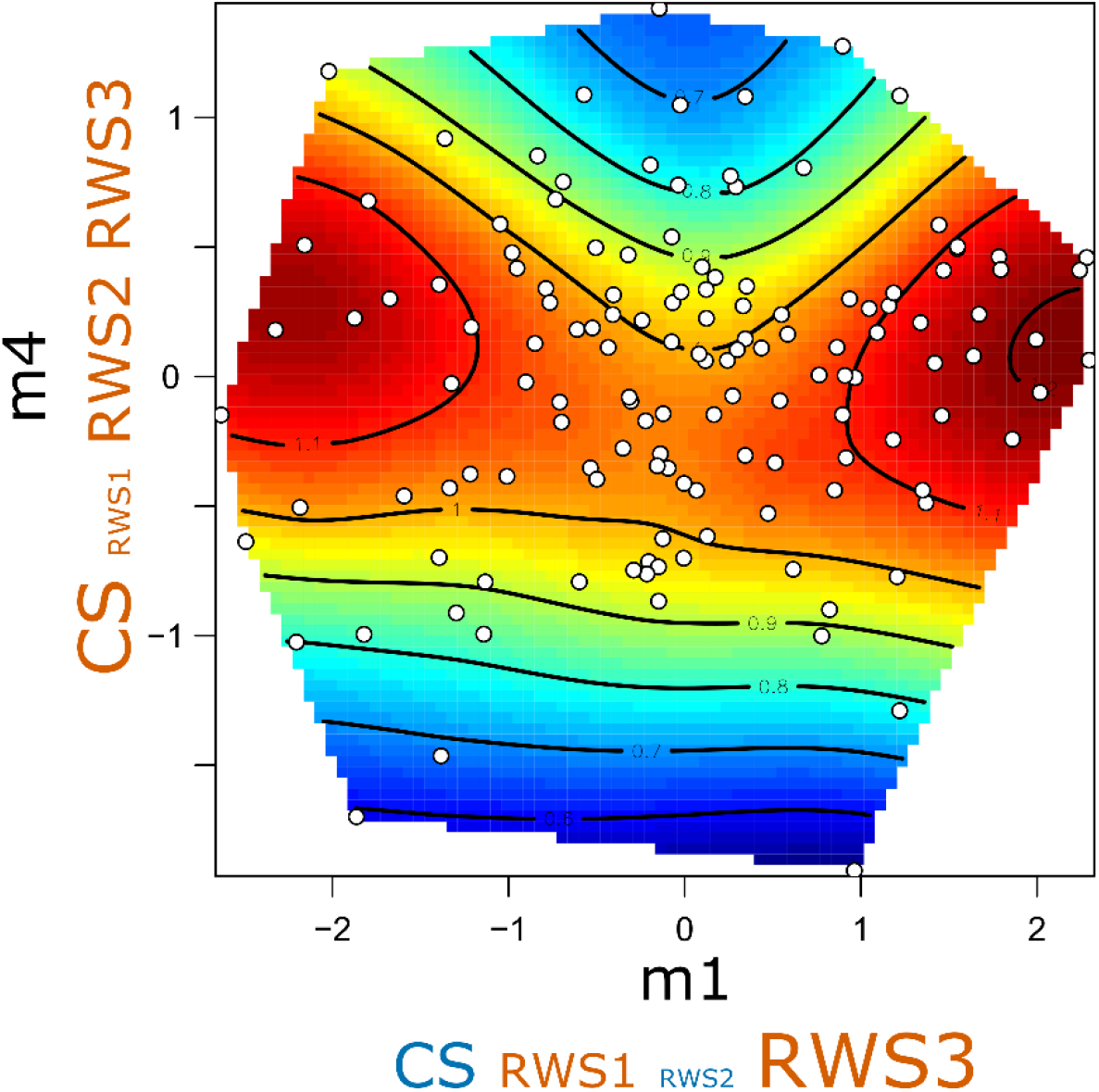
The fitness surface of the composite stylet traits on sperm transfer efficiency (*STE*\*). The axes represent the two significant eigenvectors that had positive (m1) and negative (m4) eigenvalues (Table 2B). The colour of the fitness surface corresponds to the fitness value. The font sizes of the morphological traits below the eigenvector are proportional to the square root of the trait’s |eigenvalue| to represent the respective loadings of each morphological trait on the composite traits. The colour of the morphological traits indicates positive (blue labels) and negative (orange labels) eigenvalues.

### Sperm traits set

We found linear selection in the sperm traits set on *mRS*\* (Table 1A), which arose from positive selection of brush size on *mRS*\* (step 1.2; Table 2B; Figure 2E). Worms producing sperm cells with a longer brush sired on average more offspring. Patterns of linear selection were not significantly different across fitness components (Table S1A), suggesting that the effect of brush size arises from weak selection on multiple fitness components (Table 2A).

We found nonlinear selection in the original morphological traits (Table 1A) and on the composite traits on *mRS*\* (Table 1B). The quadratic terms in the γ matrix were significant for the sperm body size and shaft size, and there were significant interactions between feeler size and shaft size, and between brush size and bristle size (Table 2A). Three composite traits significantly predicted *mRS*\* (Table 2B; Figure 2F). The two composite traits with the largest |eigenvalues|, m5 and m4, were concave and mostly loaded by sperm body size and shaft size, respectively (Figure 4). The third significant composite trait, m1, was convex and seems to oppose brush size on the one side (i.e., positive loading) and bristle size, feeler size and shaft size on the other side (i.e., negative loadings). This result may suggest that alternative sperm phenotypes may be correlated with high *mRS*\* values (i.e., sperm with either a long brush, or with long bristles, feeler, and shaft).

**Figure 4.**
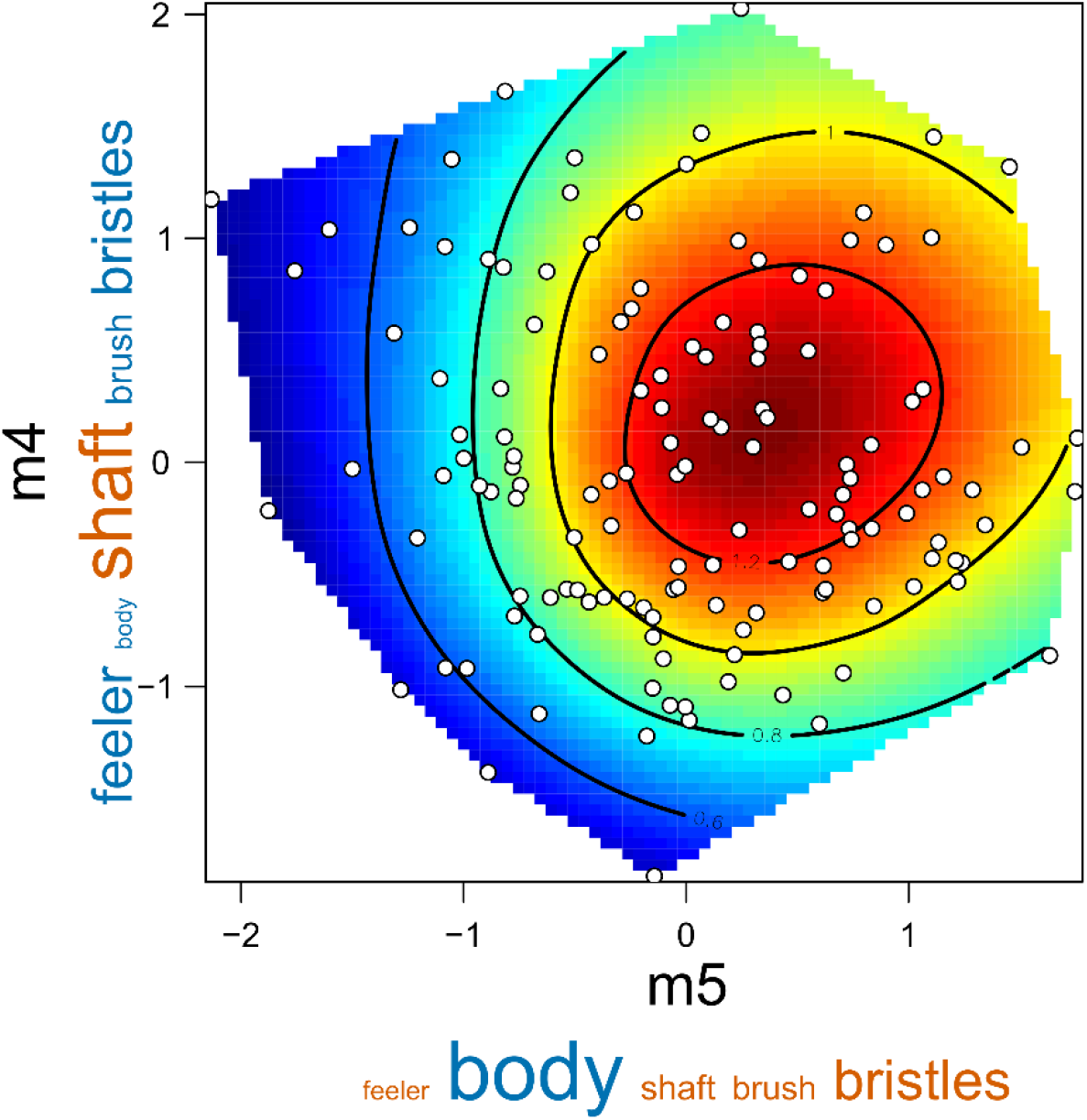
The fitness surface of the composite sperm traits on male reproductive success (*mRS*\*). The axes represent the two significant eigenvectors with the highest |eigenvalues|, m4 and m5, which were both negative (Table 2B). The colour of the fitness surface corresponds to the fitness value. The font sizes of the morphological traits below the eigenvector are proportional to the square root of the trait’s |eigenvalue| to represent the respective loadings of each morphological trait on the composite traits. The colour of the morphological traits indicates positive (blue labels) and negative (orange labels) eigenvalues.

Moreover, nonlinear selection was almost significantly different across fitness components (Table S1). We found different patterns of nonlinear selection in the sperm composite traits on *F*\* compared to *SFE*\* (Table S3B), though both fitness components induced nonlinear selection (Table 1B; Figure 2F). We found concave and convex selection on *F*\* (Table 2B). The composite trait capturing concave selection, m4, was mostly influenced by sperm body size, while the composite trait capturing convex selection, m1, was mostly influenced by feeler size and shaft size. The only significant composite trait on *SFE*\*, m3, captured concave selection and was mostly influenced by brush size (Table 2B; Figure 2F). Remarkably, other composite traits in *SFE*\* had larger eigenvalues than m3, but were not statistically significant. Altogether, our results on sperm traits showed nonlinear selection, but was spread over multiple composite traits and across fitness components, possibly rendering composite traits non-significant when considered individually.

## Discussion

Although sexual selection is acknowledged to act along multiple pre- and postcopulatory episodes of selection (Arnold and Wade 1984; Evans and Garcia-Gonzalez 2016), studies measuring and comparing phenotypic selection on pre- and postcopulatory fitness components are scarce – and even more so with respect to studies accounting for nonlinear selection through multivariate selection analyses (but see House et al. 2016). By measuring phenotypic traits alongside pre- and postcopulatory fitness components, such as mating success, sperm-transfer efficiency and sperm fertilising efficiency, we could study how the form and the strength of selection materialise along these components. Our results show that selection may arise both from specific and multiple fitness components, and that morphological traits may be under different selective pressure with respect to different fitness components. In the following, we discuss the insights gained by measuring linear and nonlinear selection on pre- and postcopulatory fitness components.

Postcopulatory selection is usually measured through a single fitness component (e.g., Devigili et al. 2015; House et al. 2016; McDonald et al. 2017; McCullough et al. 2018; De Nardo et al. 2021). Here, we could contrast phenotypic selection in two postcopulatory fitness components, sperm-transfer efficiency and sperm fertilising efficiency, finding that these two components favoured different shapes of the male copulatory organ. Namely, individuals having stylets that are curved away from the seminal vesicle (positive RWS1) and narrow (negative RWS2) had a better sperm-transfer efficiency, while sperm fertilising efficiency was higher with more bent stylet tips (negative RWS3). Although only the latter effect was significant on male reproductive success, this outcome suggests that distinct periods of selection can act on morphological traits after copulation. Our results also show nonlinear selection on stylet traits. The main source of nonlinear on stylet traits was due to an interaction between centroid size and RWS3. Selection favoured stylets that were either small and had straight tips, or large and with bent tips, which arose from sperm-transfer efficiency. In *M. lignano*, stylet shape may be selected by allowing worms to interfere with previously received sperm (e.g., sperm removal), and/or to donate sperm at strategic places in the female antrum that increase sperm fertilisation success. These two putative mechanisms may possibly induce contrasting selection on stylet shape. In our experimental design, however, because several copulations had occurred during our mating trials, it is possible that sperm abilities to resist being displaced by subsequent mating partners can translate into high sperm-transfer efficiencies.

We found positive linear selection on Δ seminal vesicle size. In *M. lignano*, the seminal vesicle is a reliable proxy for the number of sperm it contains (Schärer and Vizoso 2007), so that the increase in seminal vesicle size of worms kept in isolation for two days can serve as a proxy for the sperm production rate. Thus, this result suggest that worms with higher sperm production rates sired more offspring, which is predicted by sperm competition theory (Parker 1970; Pizzari and Parker 2009). Although, we did not find evidence for a difference in selection across fitness components, the effect of Δ seminal vesicle size was significantly positive only on mating success. This result confirms a previous study, in which experimentally manipulated sperm production rate was shown to influence both mating success and male reproductive success (Sekii et al. 2013). Worms seem to adjust their mating rates according to the amount of sperm available to donate to partners, which illustrates that pre- and postcopulatory selection can be intertwined. Therefore, studies finely decomposing sexual selection along fitness components are likely to provide critical insights into this interplay, while studies measuring phenotypic selection on total fitness are more likely to provide misleading interpretations about the biological mechanisms underlying the measured selection.

Surprisingly, we did not find selection on testis size, which clearly contrasts with several previous findings in *M. lignano*. Bigger testes have been found to positively correlate with sperm production rate (Schärer and Vizoso 2007), with the number of mating partners and the number of sperm transferred to partners (Janicke and Schärer 2009a), with sperm-transfer efficiency and male reproductive success (Marie-Orleach et al. 2016), and with paternity share (Vellnow et al. 2018). Remarkably, testis size showed one of the lowest repeatability value between the two morphological measurements taken two days apart: body size, *r_I_* = 0.66; testis size, *r_I_* = 0.36; ovary size, *r_I_* = 0.35; stylet centroid size, *r_I_* = 0.87; stylet RWS1, *r_I_* = 0.68; stylet RWS2, *r_I_* = 0.78; stylet RWS3, *r_I_* = 0.48 (Lessells and Boag 1987; Stoffel et al. 2017). In comparison, Schärer and Ladurner (2003) found a repeatability value of *r_I_* = 0.76 for testis size but, in contrast to our repeated measures, theirs were performed in quick sequence. This could suggest that, in our experiment, the worms adjusted their sex allocation within the two days of isolation (though such adjustments generally seem to take longer to materialise; Brauer et al. 2007). However, using only the first measurement of testis size did not predict male reproductive success either (selection differential±s.e.: -0.07±0.07, *t*=-0.97, *P*=0.336). It is thus unclear why, in our experiment, testis size showed low repeatability and did not correlate with male fitness.

We found that worms producing sperm cells with a longer brush sired more offspring (and we could not determine from which fitness component this effect specifically arose). It is currently unclear what are the biological functions of the sperm brush in *M. lignano*, and to our knowledge no specific hypotheses have been proposed to date. Several species within the genus *Macrostomum* lack a sperm brush, and these species reproduce through traumatic insemination (i.e., sperm donors inject ejaculate through the epidermis of the sperm recipient, and sperm cells subsequently move through the recipient body to fertilise the eggs) (Schärer et al. 2011; Brand et al. 2022). The biological functions of the sperm brush may thus possibly only be relevant when, like in *M. lignano*, mating is reciprocal, and sperm deposition and fertilisation occur inside the female antrum.

Moreover, we found evidence for nonlinear selection on sperm traits, which arose from multiple traits and multiple fitness components. We found concave selection on sperm body size and sperm shaft size, as well as convex selection on a composite trait opposing brush size to feeler size, bristle size, and shaft size (suggesting that alternative sperm phenotypes may be selected; Pizzari and Parker 2009). Our analysis suggests that this nonlinear selection on sperm traits arose from partner fecundity and sperm fertilising efficiency. However, focal effects on partner fecundity could represent a type I error, because the focals did not influence partner fecundity consistently in our experiment (Marie-Orleach et al. 2021). And, the significant composite trait on sperm fertilising efficiency did not concord much with the composite traits affecting male reproductive success. Remarkably, however, other composite traits had stronger, yet non-significant, signs of nonlinear selection on sperm fertilising efficiency. Therefore, we think that the unclear match between selection on male reproductive success and selection on its fitness components may be because sperm traits had relatively subtle effects (direct and/or indirect) on multiple fitness components, and these effects were statistically significant only when summed up all together in male reproductive success.

Finally, sexual selection is usually measured through two broadly defined approaches: (i) a trait-based approach, focusing on the relationship between variation in phenotypic traits and individual fitness, and (ii) a variance-based approach, using exclusively variance in individual success (e.g., opportunity for (sexual) selection; Arnold and Wade 1984). In our experiment, we used both approaches to study pre- and postcopulatory sexual selection (the variance-based approach being reported in Marie-Orleach et al. 2021), and thus gained complementary insights. By using the trait-based approach, we could report linear and nonlinear selection acting on pre- and postcopulatory fitness components. However, this approach is limited to the traits that can actually be measured in a study system. Unmeasured traits may also be under pre- and postcopulatory sexual selection, and could possibly change interpretations of phenotypic selection (i.e., the missing trait problem in multivariate analysis; Morrissey et al. 2012). That, for instance, explains why using trait sets is sub-optimal, as correlational selection among traits of different trait sets are not tested. In contrast, because the variance-based approach does not rely on phenotypic traits, these findings were not restricted by the traits measured, and so are possibly more informative about the total strength of selection arising from different fitness components (Marie-Orleach et al. 2021).

Moreover, we used both approaches in an experimental design where individual success was assessed in three independent groups. Such a design allowed us to assess the repeatability of individual success, and thus to treat phenotypic effects arising from partner fecundity as spurious effects because this fitness component was not repeatable over mating groups (i.e., not affected by the focal identity; Marie-Orleach et al. 2021). Going forwards, we advocate using both the variance-based and the trait-based approach, combined with repeated measures in future selection analyses. And we believe that promising research avenues will be to further include information about the genetic variance and co-variance in the phenotypic traits, and in the individual success in pre- and postcopulatory fitness components.

## Conclusions

We performed mating observations, *in vivo* sperm tracking, paternity analyses, and measured several morphological traits to study the strength and form of selection arising from pre- and postcopulatory sexual selection in the free-living flatworm *Macrostomum lignano*. We found evidence for linear selection on sperm production rate arising from multiple fitness components, and on (combinations of) stylet and sperm traits arising mostly from sperm-transfer efficiency and sperm fertilising efficiency. Our results suggest that intense periods of selection arise from different fitness components in *M. lignano*, overall inducing contrasting patterns of selection on combinations of traits. Thus, our study contributes to a better understanding of the complex interplay between pre- and postcopulatory sexual selection.

## Author contributions

LMO and LS conceived the study and designed the experiment. LMO performed the experiment and collected the data. LMO performed the statistical analyses with help from MDH. LMO wrote the manuscript with help from MDH and LS.

## Acknowledgements

We thank Christian Felber, Nikolas Vellnow, and Dita B. Vizoso for assistance during the experiment, and Jürgen Hottinger, Daniel Lüscher, and Urs Stiefel for technical support. This study was funded by a grant from the Swiss National Science Foundation to L.S. (SNF grant 31003A-143732).

## Conflict of interest

The authors declare no conflict of interest.

## Data availability

The data and scripts are in the supplementary material.

## Notes

### Competing Interest Statement

The authors have declared no competing interest.

