## Supplementary Information for "Contrasting the form and strength of pre- and postcopulatory sexual selection in a transparent worm with fluorescent sperm"

**Table S1.** Statistical outputs of a sequential model building approach (using partial *F*-test) to compare the strength and form of selection on the original traits among the four fitness components, first adding the linear terms and then also adding the nonlinear terms. Contrasting selection among fitness components is tested based on the 'by component' interaction term. Comparisons are made within each trait set. *$P$<0.05; **$P$<0.01; ***$P$<0.001.

| | General traits set | | | | | | Stylet traits set | | | | | | Sperm traits set | | | | | |
|---|---|---|---|---|---|---|---|---|---|---|---|---|---|---|---|---|---|---|
| | res df | res SS | df | SS | *F* | *P* | res df | res SS | df | SS | *F* | *P* | res df | res SS | df | SS | *F* | *P* |
| **A–Original morphological traits** | | | | | | | | | | | | | | | | | | |
| intercept | 549 | 134.3 | | | | | 549 | 134.3 | | | | | 549 | 134.3 | | | | |
| + linear | 545 | 132.2 | 4 | 2.0 | 2.03 | 0.090 | 545 | 130.5 | 4 | 3.8 | 4.24 | **0.002** | 544 | 132.1 | 5 | 2.2 | 1.75 | 0.122 |
| + linear x component | 533 | 130.0 | 12 | 2.2 | 0.74 | 0.717 | 533 | 120.7 | 12 | 9.8 | 3.65 | **<0.001** | 529 | 129.5 | 15 | 2.7 | 0.72 | 0.766 |
| + non-linear | 523 | 128.2 | 10 | 1.9 | 0.75 | 0.676 | 523 | 116.9 | 10 | 3.8 | 1.73 | 0.072 | 514 | 124.3 | 15 | 5.2 | 1.41 | 0.139 |
| + non-linear x component | 493 | 123.1 | 30 | 5.1 | 0.68 | 0.901 | 493 | 109.8 | 30 | 7.1 | 1.06 | 0.380 | 469 | 115.5 | 45 | 8.8 | 0.79 | 0.831 |
| **B–Composite traits** | | | | | | | | | | | | | | | | | | |
| intercept | 549 | 134.3 | | | | | 549 | 134.3 | | | | | 549 | 134.3 | | | | |
| + linear | 545 | 132.2 | 4 | 2.0 | 2.09 | 0.081 | 545 | 130.5 | 4 | 3.7 | 4.38 | **0.002** | 544 | 132.1 | 5 | 2.2 | 1.85 | 0.101 |
| + linear x component | 533 | 130.0 | 12 | 2.2 | 0.76 | 0.693 | 533 | 120.7 | 12 | 9.8 | 3.77 | **<0.001** | 529 | 129.5 | 15 | 2.7 | 0.76 | 0.722 |
| + non-linear | 529 | 128.2 | 4 | 1.9 | 1.94 | 0.103 | 529 | 116.9 | 4 | 3.8 | 4.46 | **0.002** | 524 | 124.3 | 5 | 5.2 | 4.47 | **<0.001** |
| + non-linear x component | 517 | 125.0 | 12 | 3.1 | 1.08 | 0.375 | 517 | 111.4 | 12 | 5.5 | 2.11 | **0.015** | 509 | 118.5 | 15 | 5.7 | 1.65 | 0.059 |

**Table S2.** Statistical outputs of a sequential model building approach (using partial *F*-test) to compare the strength and form of selection on the original traits (A) and the composite traits (B) in a pair-wise fashion (*i.e.*, between each pair of the four fitness components). Contrasting selection is tested based on the 'by component' interaction term. Comparisons only when selection was inconsistent over all fitness components (see Table S3 A). **P*<0.05; ***P*<0.01; ****P*<0.001.

**A–Original morphological traits**

*Stylet traits set*

| | | | ***MS**** | | | | ***STE**** | | | | ***SFE**** | | | |
|---|---|---|---|---|---|---|---|---|---|---|---|---|---|---|
| | | df | res. SS | SS | *F* | *P* | res. SS | SS | *F* | *P* | res. SS | SS | *F* | *P* |
| | intercept | | 23.2 | | | | 46.5 | | | | 84.7 | | | |
| ***F**** | + linear | 4 | 22.7 | 0.5 | 1.54 | 0.190 | 45.7 | 0.8 | 1.44 | 0.222 | 80.7 | 3.95 | 3.42 | **0.010**** |
| | + linear x component | 4 | 21.9 | 0.7 | 2.22 | 0.067 | 42.5 | 3.15 | 5.36 | **<0.001***** | 75.3 | 5.4 | 4.67 | **0.001*** |
| | intercept | | | | | | 49.6 | | | | 87.7 | | | |
| ***MS**** | + linear | 4 | | | | | 47.7 | 1.9 | 2.98 | **0.020*** | 81.1 | 6.7 | 5.57 | **<0.001***** |
| | + linear x component | 4 | | | | | 45.4 | 2.3 | 3.66 | **0.006**** | 78.2 | 2.9 | 2.39 | 0.051 |
| | intercept | | | | | | | | | | 111.1 | | | |
| ***STE**** | + linear | 4 | | | | | | | | | 103.9 | 7.2 | 4.93 | **0.001***** |
| | + linear x component | 4 | | | | | | | | | 98.8 | 5.1 | 3.49 | **0.009**** |

**Table S2. Continued.**

### B–Composite traits

#### *Stylet traits set*

| | | df | *MS** res. SS | SS | *F* | *P* | *STE** res. SS | SS | *F* | *P* | *SFE** res. SS | SS | *F* | *P* |
|---|---|---|---|---|---|---|---|---|---|---|---|---|---|---|
| ***F**** | intercept | | 23.2 | | | | 46.5 | | | | 84.7 | | | |
| | + linear | 4 | 22.7 | 0.5 | 1.58 | 0.180 | 45.7 | 0.8 | 1.46 | 0.214 | 80.7 | 4.0 | 3.54 | **0.008**** |
| | + linear x component | 4 | 21.9 | 0.7 | 2.27 | 0.062 | 42.5 | 3.1 | 5.46 | **0.000***** | 75.3 | 5.4 | 4.84 | **0.001***** |
| | + quadratic | 4 | 21.1 | 0.8 | 2.59 | **0.037*** | 40.1 | 2.4 | 4.23 | **0.002**** | 73.1 | 2.3 | 2.02 | 0.092 |
| | + quadratic x component | 4 | 20.9 | 0.2 | 0.75 | 0.561 | 37.5 | 2.6 | 4.52 | **0.002**** | 71.6 | 1.4 | 1.30 | 0.271 |
| ***MS**** | intercept | | | | | | 49.6 | | | | 87.7 | | | |
| | + linear | 4 | | | | | 47.7 | 1.9 | 3.07 | **0.017*** | 81.1 | 6.7 | 5.81 | **0.000***** |
| | + linear x component | 4 | | | | | 45.4 | 2.3 | 3.78 | **0.005**** | 78.2 | 2.9 | 2.50 | **0.043*** |
| | + quadratic | 4 | | | | | 42.0 | 3.4 | 5.56 | **0.000***** | 75.0 | 3.1 | 2.74 | **0.029*** |
| | + quadratic x component | 4 | | | | | 39.7 | 2.3 | 3.76 | **0.005**** | 73.8 | 1.2 | 1.06 | 0.378 |
| ***STE**** | intercept | | | | | | | | | | 111.1 | | | |
| | + linear | 4 | | | | | | | | | 103.9 | 7.2 | 5.10 | **0.001***** |
| | + linear x component | 4 | | | | | | | | | 98.8 | 5.1 | 3.61 | **0.007**** |
| | + quadratic | 4 | | | | | | | | | 92.1 | 6.7 | 4.77 | **0.001***** |
| | + quadratic x component | 4 | | | | | | | | | 90.6 | 1.5 | 1.03 | 0.390 |

#### *Sperm traits set*

| | | df | *MS** res. SS | SS | *F* | *P* | *STE** res. SS | SS | *F* | *P* | *SFE** res. SS | SS | *F* | *P* |
|---|---|---|---|---|---|---|---|---|---|---|---|---|---|---|
| ***F**** | intercept | | 23.2 | | | | 46.5 | | | | 84.7 | | | |
| | + linear | 5 | 22.7 | 0.4 | 1.14 | 0.340 | 45.0 | 1.6 | 1.99 | 0.081 | 84.2 | 0.5 | 0.36 | 0.876 |
| | + linear x component | 5 | 22.0 | 0.7 | 1.77 | 0.120 | 43.0 | 1.9 | 2.45 | **0.034*** | 83.4 | 0.8 | 0.56 | 0.731 |
| | + quadratic | 5 | 20.9 | 1.2 | 2.94 | **0.013*** | 41.2 | 1.8 | 2.24 | 0.051 | 77.7 | 5.6 | 3.84 | **0.002**** |
| | + quadratic x component | 5 | 20.1 | 0.8 | 1.91 | 0.093 | 40.5 | 0.7 | 0.90 | 0.482 | 74.0 | 3.7 | 2.54 | **0.029*** |
| ***MS**** | intercept | | | | | | 49.6 | | | | 87.7 | | | |
| | + linear | 5 | | | | | 47.2 | 2.4 | 2.79 | **0.018*** | 86.6 | 1.1 | 0.71 | 0.618 |
| | + linear x component | 5 | | | | | 46.1 | 1.1 | 1.32 | 0.258 | 86.4 | 0.2 | 0.14 | 0.983 |
| | + quadratic | 5 | | | | | 43.6 | 2.5 | 2.93 | **0.014*** | 80.1 | 6.3 | 4.15 | **0.001**** |
| | + quadratic x component | 5 | | | | | 43.3 | 0.4 | 0.44 | 0.821 | 77.3 | 2.8 | 1.81 | 0.111 |
| ***STE**** | intercept | | | | | | | | | | 111.1 | | | |
| | + linear | 5 | | | | | | | | | 107.9 | 3.2 | 1.62 | 0.154 |
| | + linear x component | 5 | | | | | | | | | 107.4 | 0.5 | 0.26 | 0.932 |
| | + quadratic | 5 | | | | | | | | | 100.6 | 6.8 | 3.51 | **0.004**** |
| | + quadratic x component | 5 | | | | | | | | | 98.3 | 2.3 | 1.18 | 0.319 |

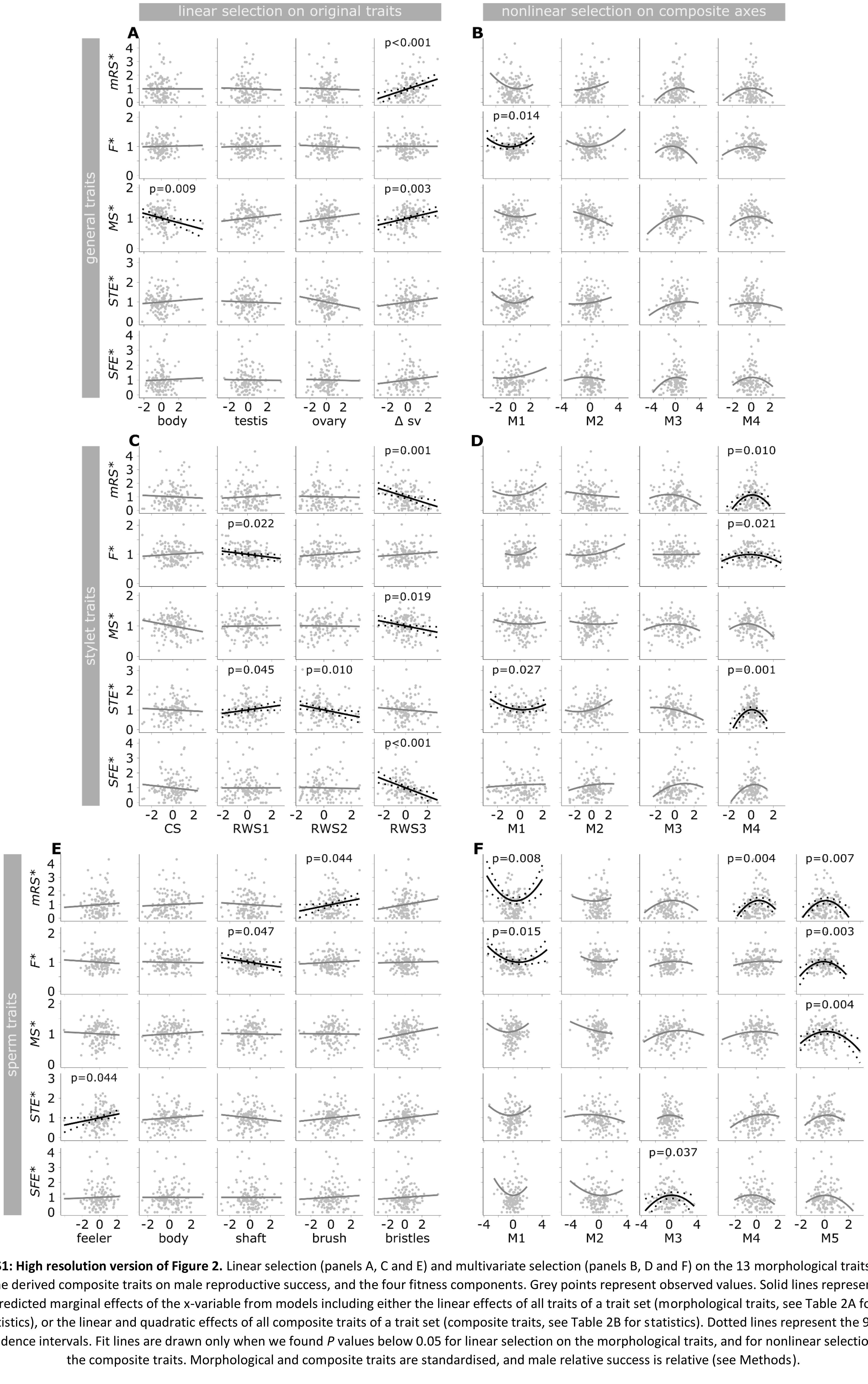


**Fig. S1: High resolution version of Figure 2.** Linear selection (panels A, C and E) and multivariate selection (panels B, D and F) on the 13 morphological traits and the derived composite traits on male reproductive success, and the four fitness components. Grey points represent observed values. Solid lines represent predicted marginal effects of the x-variable from models including either the linear effects of all traits of a trait set (morphological traits, see Table 2A for statistics), or the linear and quadratic effects of all composite traits of a trait set (composite traits, see Table 2B for statistics). Dotted lines represent the 95% confidence intervals. Fit lines are drawn only when we found *P* values below 0.05 for linear selection on the morphological traits, and for nonlinear selection on the composite traits. Morphological and composite traits are standardised, and male relative success is relative (see Methods).
